# Manner of death and demographic effects on microbial community composition in organs of the human cadaver

**DOI:** 10.1101/752576

**Authors:** Holly Lutz, Alexandria Vangelatos, Neil Gottel, Emily Speed, Antonio Osculati, Silvia Visona, Sheree J. Finley, Sari Tuomisto, Pekka Karhunen, Jack A. Gilbert, Gulnaz T. Javan

## Abstract

The microbiome serves important functions in human health, and postmortem, the microbial signatures of colonized organ tissue could be useful in helping to predict the manner of death in cases where this information is not known. We surveyed the microbiota (16S rRNA V4 amplicon sequencing) of 265 organ tissue samples including liver, blood, brain, heart, prostate, spleen and uterus from cadavers in Italy, Finland and the United States with confirmed manners of death comprising either accidental death, natural death, homicide, and suicide. Geographic locality (i.e. nationality) had a strong effect on observed microbial composition. Differing PERMANOVA results between unweighted and weighted UniFrac (nearly inverse results) suggest that specific bacteria may be associated with ethnicity and age, but that these differences are negligible when taking into account the relative abundance of bacterial taxa; weighted UniFrac measures suggest that although taxonomic composition may not vary significantly between different manners of death, PMI, or BMI categories, the relative abundance of specific taxa vary significantly. Various tissues exhibit differential associations with bacteria, and prostate and uterus were substantially different compared to other organs. For example, in Italian cadavers, the bacteria MLE1-12 permeated nearly all tissues, except the prostate and uterus. We identified specific bacterial ASVs as biomarkers of either natural or accidental death and suicide, but not for homicide. While the manner of death may have an impact on microbial associations, further investigation under more controlled conditions will be needed to validate whether these associations are predictive in forensic determinations.

**Importance:** The utilization of microbial data in the context of forensic investigations holds great promise for the field of forensic science. Identification of taxa that are associated with postmortem interval (PMI), specific manners of death (MOD), or other traits such as age, sex, ethnicity, and nationality may allow investigators to refine the circumstantial details surrounding the death of an individual. In this study we find nationality (geographic location of cadaver) to be a dominant predictor of cadaver microbiome composition. We also identify a number of cadaver-specific traits to be associated with microbial alpha- and beta diversity, as well as bacterial taxa that are differentially associated with these traits.

## 1. Introduction

During life, the microbiome serves important health-related functions including nutrient acquisition, pathogen defense, energy salvage, and immune defense training (1). The microbiome has also been linked to cardiovascular, metabolic and immune disease, as well as mental health disorders via the gut-brain-axis (2). Upon death, microbial communities present within and on the body are exposed to radical environmental changes, and recent studies have shown that microbial succession among mammalian cadavers follows a metabolically predictable progression (3, 4).

Forensic microbiology represents a potential emerging discipline in which microorganisms serve as forensic tools or trace evidence. Advances in DNA sequencing technologies paired with increased understanding of the human microbiome have hinted at the possibility that the microbiome could be used as a biomarker of decay (3) and as trace evidence to link individual people to objects they have previously interacted with (5–9). Recent studies have also shown that the microbiome can be used to estimate the amount of time that has elapsed since death, referred to as the postmortem interval (PMI), allowing investigators to establish a potential timeline of death (3, 10–16).

The microbial composition and abundance associated with internal organ tissues are dependent on temperature, manner/cause of death, and PMI, since bacteria have different growth optima based on the physicochemical constraints of their environment (17–19). Also microbial abundance associated with the body antemortem can play a role in decay, as a cadaver of an aged adult human, with approximately 40 trillion microbial cells, decays more rapidly than a deceased fetus or newborn, which usually have reduced microbial colonization density (20). Of course, these trends are contingent upon the medications and disease state of the individual.

Here we investigate the extent to which microbial associations among different organs in human cadaver can be used to predict manner of death (MOD), PMI, and geographic locality of origin. By sampling human cadavers from three disparate geographic origins (Finland, Italy, and the United States), we were able to ascertain that geographic locality has a significant influence on microbial community composition of postmortem tissues, and that despite these differences, commonalities may still be identified both among tissues, and individuals who died due to varying causes of death (e.g. natural, accidental, homicidal, and suicidal deaths). We were unable to detect significant correlations between various samples and the postmortem interval, likely due to the fact that the sampling regimen was optimized to capture variation among geographic locality, organ type, and manner of death. Significant patterns were observed in this study associated with geography and manner of death warrant reinforcement from additional investigations to elucidate the origin of these associations.

## 2. Methods

### 2.1 Sampling

Postmortem samples included corpses from the Alabama Department of Forensic Sciences in Montgomery, AL, USA and The Office of the District One Medical Examiner in Pensacola, FL, USA; Pavia University in Italy; and Tampere University in Finland. Demographic data were collected on each of the corpses. Corpses were kept in a morgue at 4°C until the time of tissue collection. The age, sex, BMI, height, ethnicity, and PMI were documented for each corpse. Tissue sampling was performed in an examination area with an ambient temperature of 20°C. Sections of the internal organs was dissected using a sterile scalpel and placed in labeled, sterile polyethylene bags. For the USA samples, tissues were transported from the morgue to the laboratory on ice and immediately frozen at −80°C until processing. DNA was extracted from internal organs by conventional chemical and physical disruption protocols (21) using the phenol chloroform method, which is specifically optimized for recovery of microbial DNA from low-yield samples. The quality and quantity of DNA was determined by spectrophotometry (NanoDrop™).

### 2.2 DNA extraction and sequencing and statistical analyses

We used the standard 515F and 806R primers (22–24) to amplify the V4 region of the 16S rRNA gene, using mitochondrial blockers to reduce amplification of host mitochondrial DNA. Sequencing was performed using paired-end 150 base reads on an Illumina HiSeq sequencing platform. Following standard demultiplexing and quality filtering using the Quantitative Insights Into Microbial Ecology pipeline (QIIME2) (25) and vsearch8.1 (26), Absolute Sequence Variants (ASVs) were identified using the Deblur method (27) and taxonomy was assigned using the Greengenes Database (May 2013 release; http://greengenes.lbl.gov).

### 2.3 Statistical analyses

Following quality filtering and taxonomy assignment, sequence libraries were rarefied to a read depth of 5,000 reads, and rarefied libraries were used for all subsequent analyses. Alpha diversity was calculated using the Shannon index, and measured species richness based on actual observed diversity. Significance of differing mean values for each diversity calculation was determined using the Kruskal-Wallis rank sum test, followed by a post-hoc Dunn test with Benjamini-Hochberg corrected *p*-values. Two measures of beta diversity (unweighted UniFrac and weighted UniFrac) were calculated using relative abundances of each ASV (calculated as ASV read depth divided by total library read depth). Significant drivers of community similarity were identified using the ADONIS test with Bonferroni correction for multiple comparisons using the R package Phyloseq (28). ANCOM analyses were performed to assess significance of differential abundance based on log2fold change measures between categories (e.g. geographic localities, organ types, and manners of death). Analyses were run independently for each variable, e.g. ASVs associated with Finnish cadavers were compared to ASVs from all other localities grouped together, ASVs associated with Italian cadavers were compared to those from all other localities, and so on. Additional R packages used for analyses and figure generation included vegan (29), ggplot2 (30), and dplyr (31). For a complete list of packages and codes for microbiome analyses, see http://github.com/hollylutz/CadaverMP. All 16S rRNA sequence and sample metadata are publicly available via the QIITA platform under Study ID #### and the European Bioinformatics Institute (EBI) under accession number ####.

## 3. Results

### 3.1 Cadaver and organ sampling and analysis

We collected 265 samples of multiple organs from corpses derived from Finland, Italy, and the United States (Table S1). Sampling spanned PMIs of 3.5 to 432 hours (avg = 87.6 hours) and included tissues from cadavers corresponding to different manners of death grouped into four categories: accidental death (n = 88), natural death (n = 106), homicide (n = 23), and suicide (n = 45) (Table 2). In total, 4,337,301 16S rRNA V4 amplicon sequencing reads were generated from 265 samples, comprising 2,204 ASVs. Following sequence deblurring and rarefaction analysis (5,000 read per library cut-off), we identified 1,855 ASVs across 163 remaining samples (Table 1), with a range of 239 to 1,413 ASVs across different organs.

**Table 1.** Sampling by geographic location and organ, post-rarefaction

| Location | Organ |  |  |  |  |  |  |
| --- | --- | --- | --- | --- | --- | --- | --- |
|  | Blood | Brain | Heart | Liver | Prostate | Spleen | Uterus |
| Finland | 0 | 0 | 0 | 20 | 0 | 0 | 0 |
| Italy | 0 | 10 | 9 | 13 | 13 | 10 | 5 |
| USA | 6 | 9 | 11 | 47 | 0 | 10 | 0 |
| **Total** | **6** | **19** | **20** | **80** | **13** | **20** | **5** |

**Table 2.** Sampling by manner of death and geographic locality, post-rarefaction, with PMI statistics; undetermined MOD (n=2) excluded from analyses.

| Location | Manner of Death |  |  |  | Postmortem Interval |  |  |  |
| --- | --- | --- | --- | --- | --- | --- | --- | --- |
|  | Accident (n) | Natural (n) | Homicide (n) | Suicide (n) | PMI<sub>min</sub> (hrs) | PMI<sub>max</sub> (hrs) | PMI<sub>avg</sub> (hrs) | PMI<sub>SD</sub> (hrs) |
| Finland | 6 | 12 | 0 | 2 | 48 | 192 | 108 | 42.3 |
| Italy | 20 | 24 | 3 | 13 | 24 | 432 | 112 | 96.4 |
| USA | 31 | 29 | 9 | 12 | 3.5 | 240 | 37.8 | 47.8 |
| **Total** | **57** | **65** | **12** | **27** |  |  |  |  |

### 3.2 Alpha diversity

Alpha diversity, calculated as observed number (richness) of ASVs and the Shannon diversity index, differed significantly between some but not all organs and varied by locality (both, *p* < 0.05, Kruskal-Wallis). Post-hoc tests (corrected for multiple comparisons using the Benjamini-Hochberg method) revealed that among Italian subjects, the prostate and uterus differed significantly from all other organs (brain, heart, liver, and spleen) in both observed richness (*p* < 0.05, Dunn’s Test) and Shannon diversity (*p* < 0.05, Dunn’s Test), but they did not differ significantly from each other (Fig. 1A and 1B). Among subjects from the United States (USA), the only organs that differed significantly by Shannon diversity were heart and liver (*p* = 0.032, Dunn’s Test), and no organs differed significantly by observed richness (Fig.1A). A comparison of alpha diversity measures for liver samples from all three localities (Finland, Italy, USA) identified significant differences in both observed richness and Shannon diversity between liver tissue from Finland and the USA (*p* < 0.05, Dunn’s Test), and Finland and Italy (*p* < 0.05, Dunn’s Test), but not between Italian and US livers (Fig. 1C).

**Figure 1.**
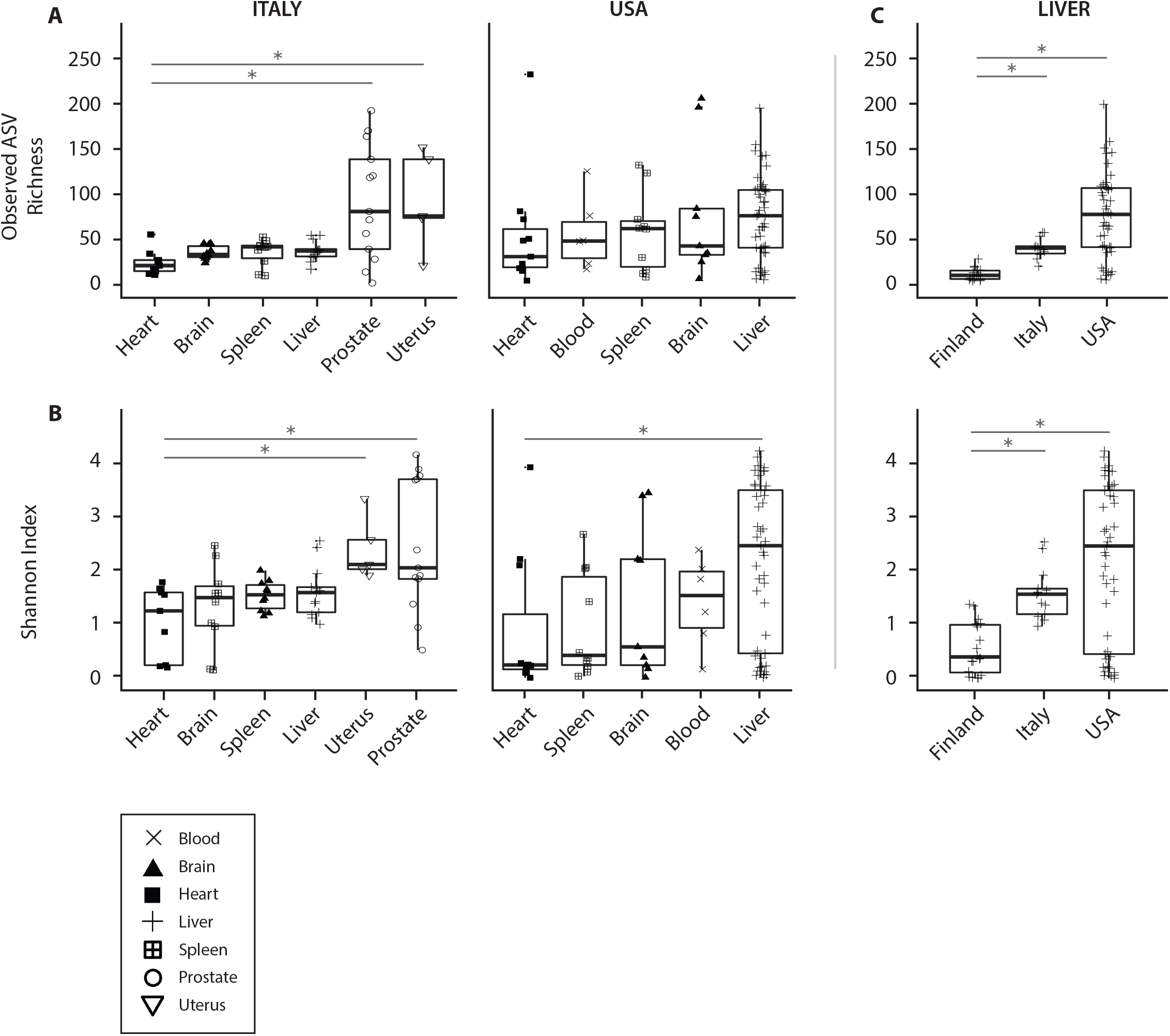
Variation in alpha diversity by organ type, A) comparing observed ASV richness between organs from different localities, B) comparing Shannon diversity index between organs from different localities. Asterisks indicate significant difference between groups based on post-hoc Dunn’s Tests, *p* < 0.05.

Alpha diversity differed significantly by manner of death among USA organs (*p* < 0.05, Kruskal-Wallis), but not among Italian or Finnish organs. Among USA organs, observed richness differed significantly between accidental deaths and homicides (*p* < 0.0177, Dunn’s Test), accidental deaths and suicides (*p* < 0.0002, Dunn’s Test), and natural deaths and suicides (*p* < 0.0005, Dunn’s Test), but not between homicides and suicides, natural deaths and accidents, or natural deaths and homicides (Fig. 2A). Also among USA organs, Shannon diversity differed significantly between accidental deaths and suicides (*p* < 0.0001, Dunn’s Test), natural deaths and homicides (*p* < 0.008, Dunn’s Test), and natural deaths and suicides (*p* = 0.000, Dunn’s Test), but not between homicides and suicides, natural deaths and accidents, or accidental deaths and homicides (Fig. 2B).

**Figure 2.**
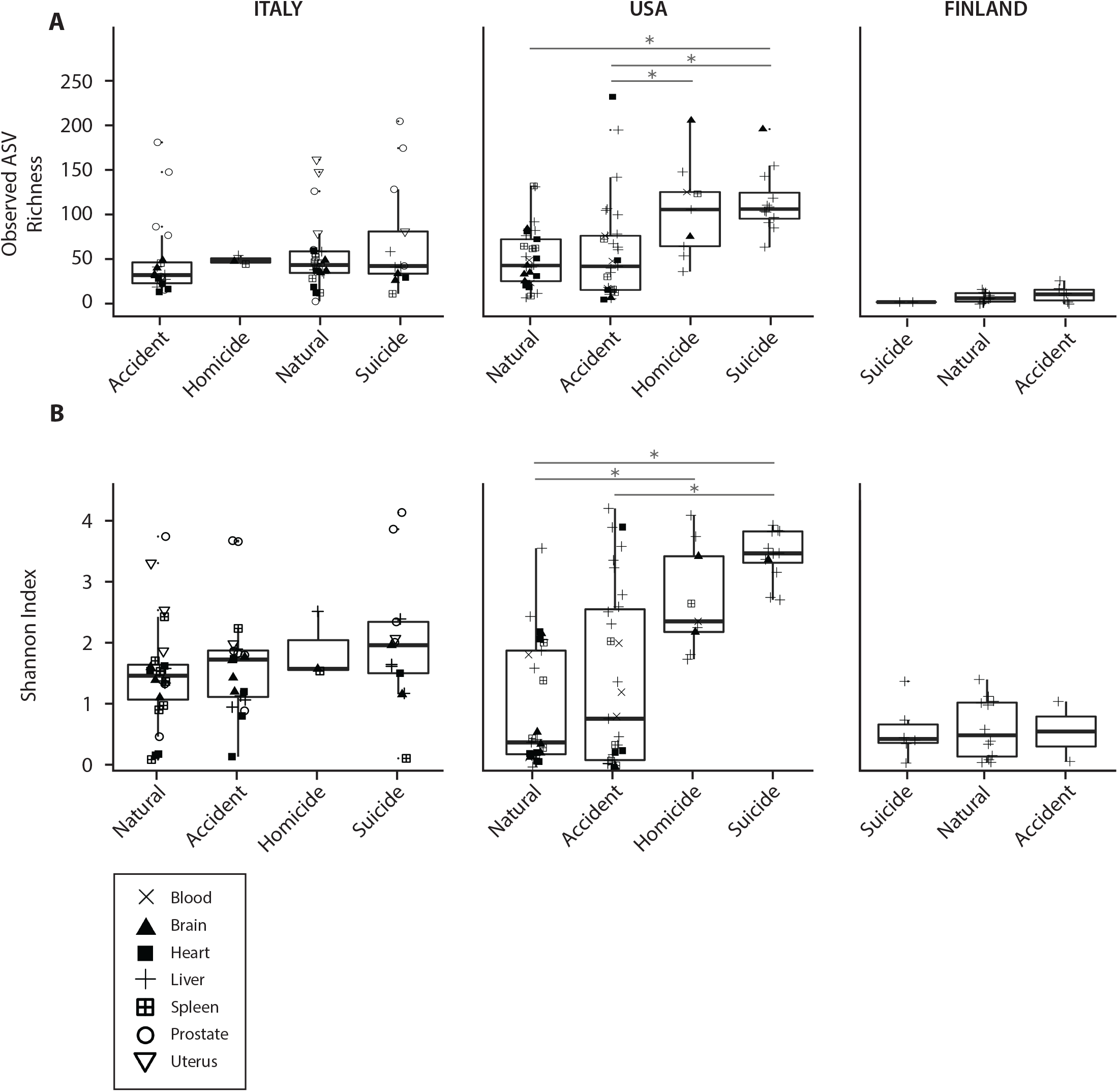
Variation in alpha diversity by manner of death, A) comparing observed ASV richness between manners of death from different localities, B) comparing Shannon diversity index between manners of death from different localities. Asterisks indicate significant difference between groups based on post-hoc Dunn’s Tests, *p* < 0.05.

Using linear regression of alpha diversity against PMI, the only significant associations observed were among Italian spleens (observed richness: *p* = 0.016, R^2^ = 0.48; Figure S1) and Finnish livers (Shannon Index: *p* = 0.021, R^2^ = 0.22; Figure S2). Similarly, we found little evidence for a correlation between BMI and bacterial alpha diversity among organs, with the exception of the Italian prostate (observed richness: *p* = 0.019, R^2^ = 0.35; Figure S3; Shannon Index: *p* = 3.58 e −05, R^2^ = 0.62; Figure S4) and the US spleen (Shannon Index: *p* = 0.017, R^2^ = 0.47; Figure S4).

### 3.3 Beta diversity

Analysis of beta diversity, using unweighted UniFrac, found a strong effect of geographic locality on postmortem bacterial community composition (Fig. 3A), whereby the microbial composition and compositional proportion were significantly different between each country (PERMANOVA: unweighted UniFrac, *p* = 0.001, R^2^ = 0.18; weighted UniFrac, *p* = 0.001, R^2^ = 0.12). No clear differences in beta diversity were visible by organ type (Fig. 3B) or organ type within each country, except for the uterus and prostate differing from all other organs in Italy (Fig. 3C), although, organ was technically a significant predictor of beta diversity (PERMANOVA: unweighted UniFrac, *p* = 0.001, R^2^ = 0.08; weighted UniFrac, *p* = 0.001, R^2^ = 0.06). Controlling for locality as a confounding variable, PERMANOVA analyses of weighted and unweighted UniFrac diversity metrics identified a number of variables significantly associated with microbial beta diversity, though these variables differed between the two metrics (Table 3). For unweighted UniFrac, significant variables included ethnicity and age (*p* < 0.05, PERMANOVA). For weighted UniFrac, significant variables included manner of death, PMI, and BMI (*p* < 0.05, PERMANOVA).

**Table 3.** PERMANOVA analysis assessing marginal effects of variables on weighted and unweighted UniFrac beta diversity, controlling for geographic locality (ADONIS, strata = locality); asterisk indicates Bonferroni adjust *p*-value < 0.05.

| | Df | SumOfSqs | R<sup>2</sup> | F | $p$-value |
| --- | --- | --- | --- | --- | --- |
| Unweighted UniFrac |  |  |  |  |  |
| Sex | 1 | 0.41 | 0.01 | 1.7 | 0.36 |
| Ethnicity | 3 | 4.03 | 0.09 | 5.46 | 0.006* |
| Age | - | 0.77 | 0.02 | 2.95 | 0.012* |
| Manner of death | 3 | 1.2 | 0.03 | 1.57 | 0.096 |
| PMI | - | 0.37 | 0.01 | 1.49 | 0.56 |
| BMI | - | 0.41 | 0.01 | 1.65 | 0.26 |
| Weighted UniFrac |  |  |  |  |  |
| Organ | 6 |  |  |  |  |
| Sex | 1 | 0.07 | 0.01 | 2.32 | 0.35 |
| Ethnicity | 3 | 0.37 | 0.06 | 3.85 | 0.73 |
| Age | - | 0.1 | 0.02 | 3.23 | 0.14 |
| Manner of death | 3 | 0.23 | 0.04 | 2.33 | 0.006* |
| PMI | - | 0.11 | 0.02 | 3.41 | 0.018* |
| BMI | - | 0.14 | 0.02 | 4.21 | 0.006* |

**Figure 3.**
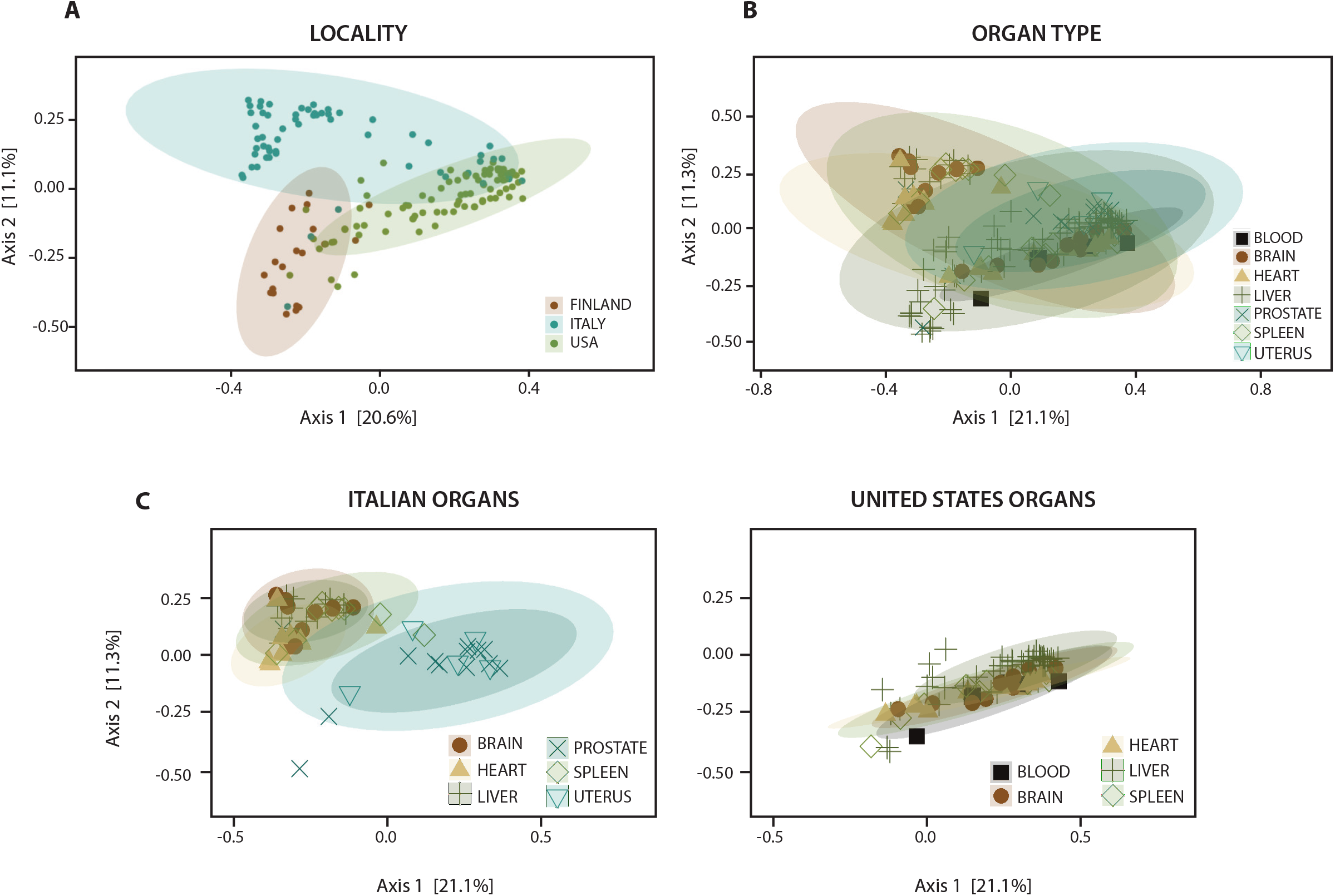
PCoA plots of unweighted UniFrac beta diversity, A) labeled by geographic locality, B) labeled by organ, and C) labeled by organ and faceted by geographic locality (Finland not included, as only liver was sampled).

### 3.4 Specific bacterial taxa associated with different factors

Analysis of composition of microbiomes (ANCOM) between different localities, organs, and manners of death identified significant differences in relative abundance (measured as the log2fold change in 16S rRNA ASV read counts) of multiple bacterial taxa. Assessing differences between localities (controlling for age, sex, BMI, PMI, ethnicity, and organ), we found that Finnish cadavers exhibited enrichment of two ASVs in the class Bacilli, as well ASVs belonging to the Alphaproteobacteria and Gammaproteobacteria, relative to cadavers from Italy and the United States (p < 0.05, ANCOM). Among Italian cadavers, we observed enrichment for ASVs in the classes Saprospirae, 4C0d-2 (phylum Cyanobacteria), Betaproteobacteria, Gammaproteobacteria, and Gemmatimonadetes (*p* < 0.05, ANCOM). And among US cadavers, we observed a significant enrichment in ASVs annotated to the class Clostridia, as well as enrichment of several ASVs belonging to the classes Alphaproteobacteria, Bacilli and Bacteroidia (p < 0.05, ANCOM) (Fig. 4; Table S2).

**Figure 4.**
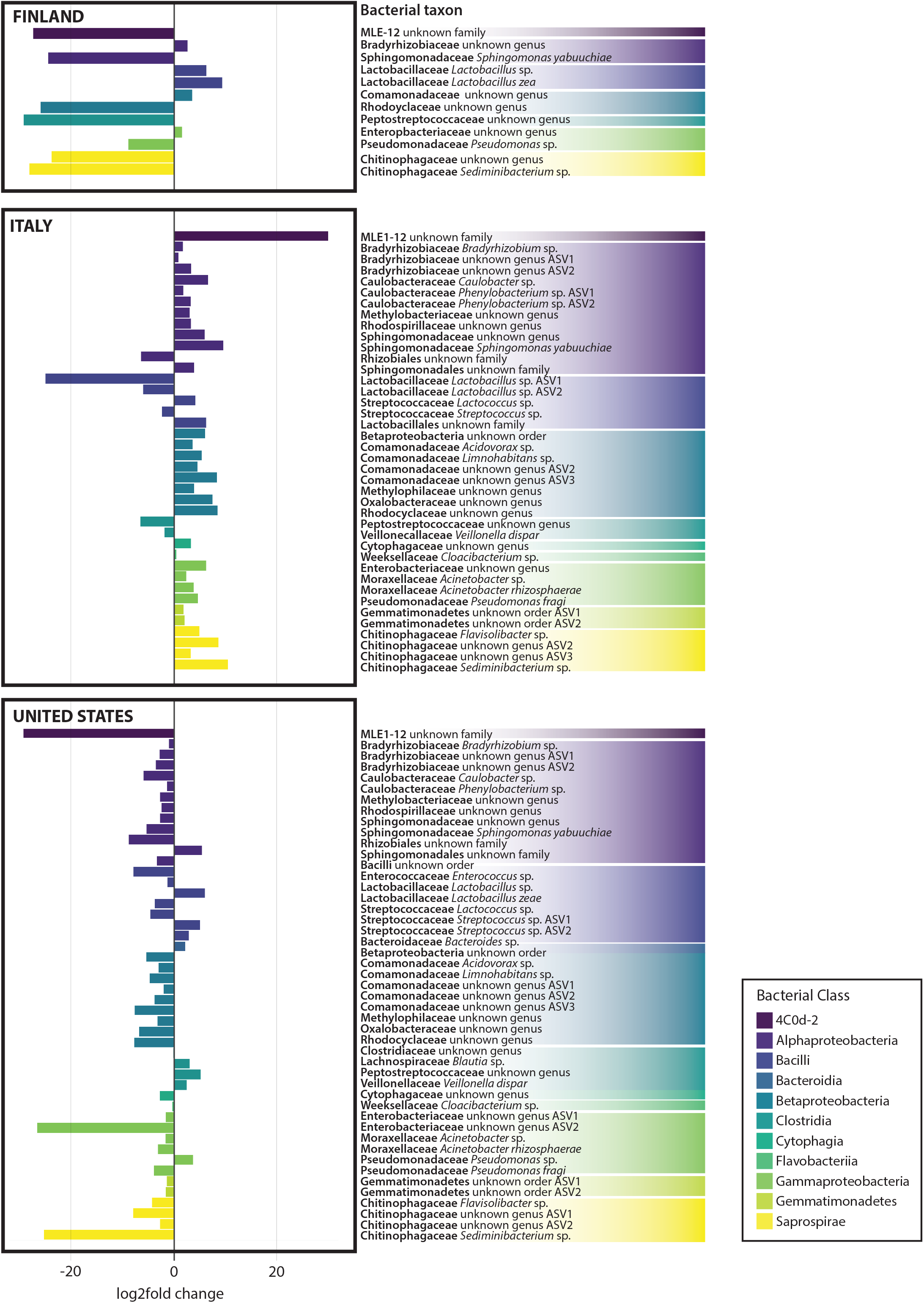
ANCOM – log2fold change in relative abundance between different cadaver localities, controlling for age, sex, ethnicity, BMI, PMI, and organ as covariates. ASVs are colored by bacterial class.

Analysis of differences in bacterial relative abundance between organs (controlling for age, sex, BMI, PMI, ethnicity, and locality) found increased proportion of a single Clostridia ASV (family Peptostreptococcaceae) in the blood, and a single Gammaproteobacteria in the heart (family Pseudomonadaceae, *Pseudomonas* sp.). Among brain tissue, a number of bacterial taxa were found to be underrepresented relative to all other organs, and none were found to be significantly enriched. Both liver and spleen exhibited an increased relative abundance of a bacterial ASVs in the class 4C0d-2 (order MLE1-12, unknown family), as well as *Sphingomonas yabuuchiae* (family Sphingomonadaceae). Other bacterial ASVs enriched in both the liver and spleen included those from classes Betaproteobacteria (specifically a single ASV in the family Rhodocyclaceae), Clostridia (specifically a single ASV in the family Peptostreptococcaceae), and Saprospirae (specifically two ASVs in the family Chitinophagaceae, and one ASV in the genus *Sediminibacterium*). The liver and prostate were both enriched for two ASVs in the class Bacteroidia, one in the family Comamonadaceae (genus *Limnohabitans*) and another in the family Oxalobacteraceae (unknown genus). The liver alone was enriched for several bacterial taxa not seen in other organs, including a Clostridia ASV in the family Lachnospiraceae (genus *Blautia*), an Alphaproteobacteria ASV in the order Rhizobiales (unknown family), and a Gammaproteobacteria in the family Enterobacteriaceae (genus *Salmonella*). Uterine tissues were enriched for only two ASVs, which were not found to be enriched in any other organs, including a single ASV in the class Bacilli (family Lactobacillaceae, genus *Lactobacillus*) and a single ASV in the class Gammaproteobacteria (family Enterobacteriaceae, unknown genus). Lastly, among prostate tissues we found a significant underrepresentation of the same 4C0d-2 ASV (order MLE1-12) observed in both liver and spleen, and a single Clostridia ASV (family Lachnospiraceae, unknown genus) relative to all other organs (except for brain, which was also depauperate with respect to the 4C0d-2 ASV) (Fig. 5; Table S3).

**Figure 5.**
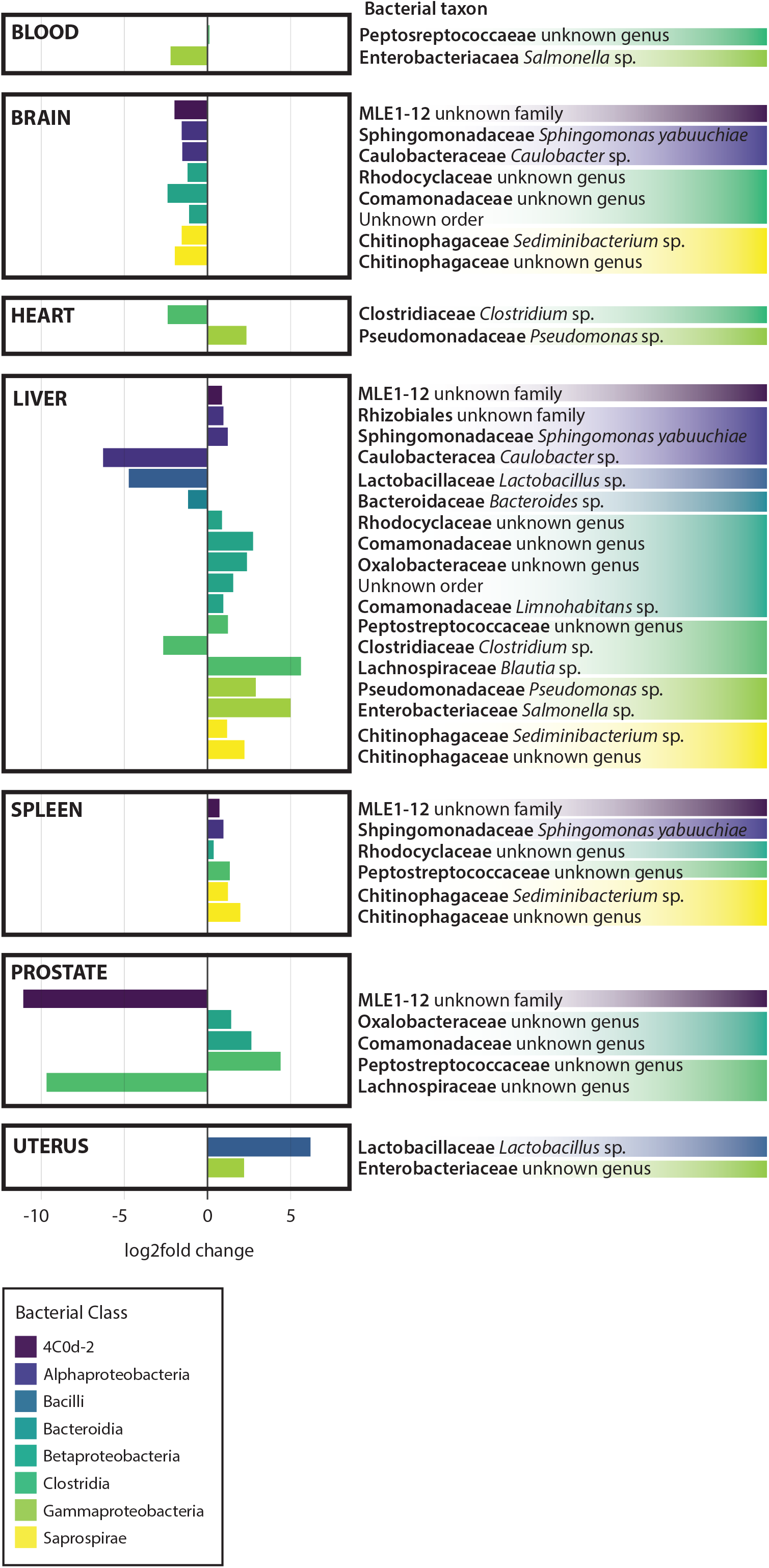
ANCOM – log2fold change in relative abundance between different organs, controlling for age, sex, ethnicity, BMI, PMI, and locality as covariates. ASVs are colored by bacterial class.

A number of unique associations between ASVs and manner of death (controlling for age, sex, BMI, PMI, ethnicity, locality, and organ) were observed. For natural deaths, this included an enrichment of the same ASV in class 4C0d-2 (order MLE1-12) mentioned previously, as well as enrichment for single ASVs in the classes Bacilli (family Lactobacillaceae, *Lactobacillus zeae*), Gammaproteobacteria (family Enterobacteriaceae, unknown genus), and Saprospirae (family Chitinophagaceae, genus *Sediminibacterium*). Among victims of accidental death, a single Bacilli ASV (order Lactobacillales, unknown family) and Gammaproteobacteria (family Enterobacteriaceae, unknown genus) were enriched. Homicide victims did not exhibit enrichment of any bacterial taxa, but exhibited a decreased abundance of ten different ASVs belonging to the class Bacilli, as well as ASVs in the classes Bacteroidia (family Prevotellaceae, *Prevotella melaninogenica*), Clostridia (family Veillonelliceae, *Veillonella dispar*), and Gammaproteobacteria (family Enterobacteriaceae, genus *Salmonella*) relative to other samples. Lastly, victims of suicide showed a similar decrease in the same Gammaproteobacteria ASV (family Enterobacteriaceae, *Salmonella*) as homicide victims, as well as decreases in another gammaproteobacterium ASV (family Pseudomonadaceae, genus *Pseudomonas*), and two ASVs in the class Clostridia (family Peptostreptococcaceae, unknown genus, and family Ruminococcaceae, *Faecalibacterium prausnitzii*). Other Clostridia ASVs were enriched in suicide victims, including two ASVs in the family Lachnospiraceae (genus *Blautia*), and one in the family Clostridiaceae (genus *Clostridium*). The only ASV belonging to class Alphaproteobacteria (order Rhizobiales) with significantly different relative abundance among manner of death categories was found to be enriched in suicide victims as well (Fig. 6; Table S4).

**Figure 6.**
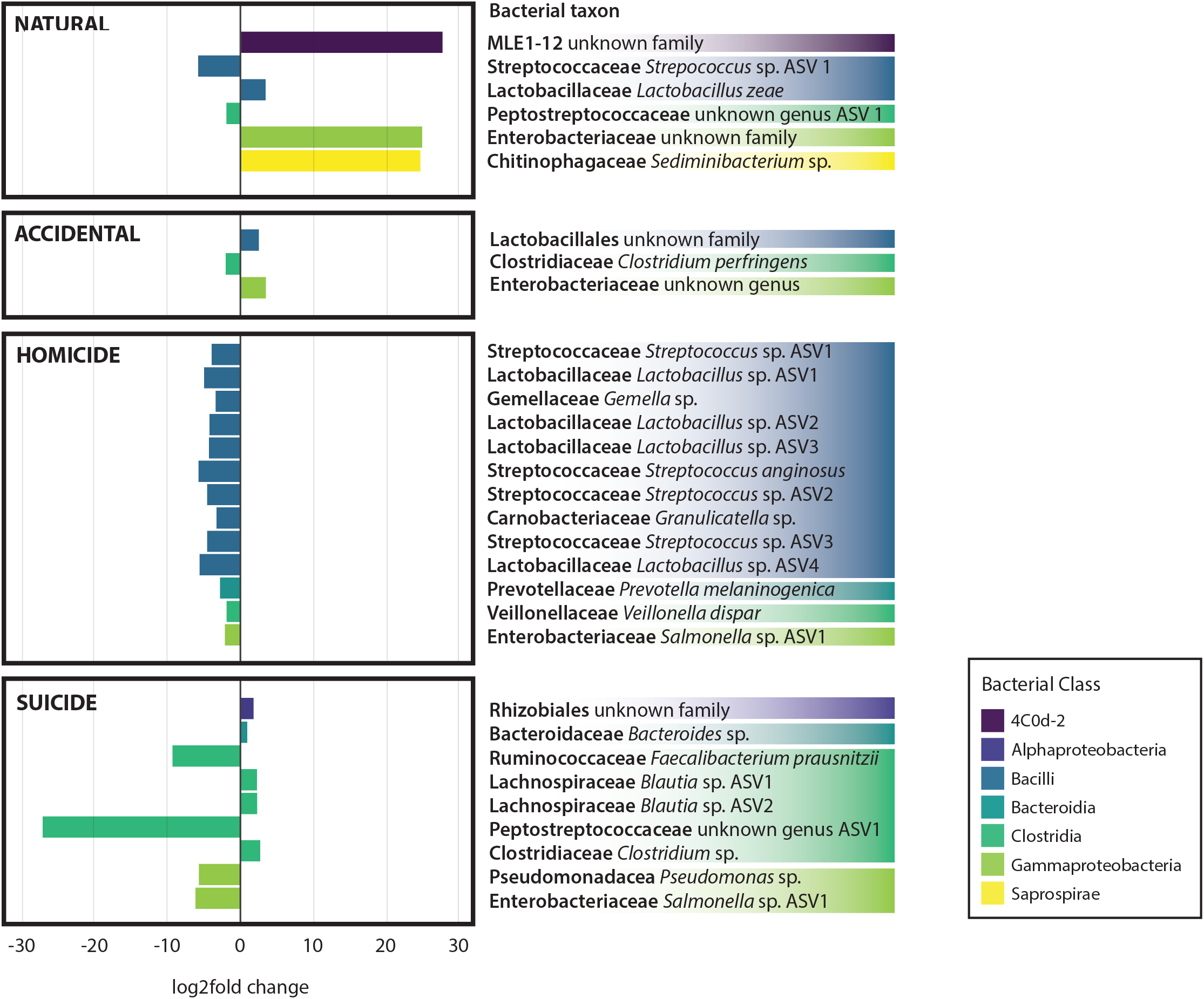
ANCOM – log2fold change in relative abundance between different manners of death, controlling for age, sex, ethnicity, BMI, PMI, organ, and locality as covariates. ASVs are colored by bacterial class.

## 4. Discussion

In this investigation we have compared existing data and newly collected samples from different organs associated with cadavers from Italy, Finland and the United States of America. We demonstrate that both the microbial alpha and beta diversity shows differential associations between organ tissue type, country or origin, and manner of death; but PMI and BMI show very few significant associations. However, the lack of consistency of the association of microbial diversity with these cadaver metrics suggests that neither alpha or beta diversity metrics would be reliable predictors of country of origin or manner of death. However, we did identify specific bacterial taxa that were enriched in differential organs, and that were significant associated with both country of origin and manner of death. This suggests potential biomarkers of manner of death could be possibly validated through further and independent experimentation, observation and validation.

A previous microbial survey of internal organ tissues (e.g., brain, heart, liver, and spleen) of four cadavers, associated with a homicide, suicide, over-dose, and accidental death cases, demonstrated that the obligate anaerobe, *Clostridium* was found in cadavers of varying PMIs, while the facultative anaerobe, *Lactobacillus*, was more abundant in cadavers with shorter PMIs (12). Other investigations performed exploratory analyses of bacteria present in mouth and rectal scrapings taken at the onset and end of the bloat stage of corpses decomposing in a natural setting (32). However, internal organs were not sampled across time points in this study. Another postmortem microbiome study of 33 bodies was conducted using bacterial culturing and reverse transcriptase quantitative PCR (RT-qPCR) techniques to profile the microbes in blood, liver, portal vein, mesenteric lymph node, and pericardial fluid, and identified 21 genera, with the most abundant being *Staphylococcus* sp., *Streptococcus* sp., *Clostridium* sp., *Enterococcus* sp., and *Escherichia* sp. (33)

We identified many different taxa as being associated with manner of death, including *Lactobacillus*, Enterobacteriaceae, *Sediminibacterium, Blautia*, Rhizobiales, and *Clostridium.* In several recent postmortem microbiome studies, the clostridia were observed to proliferate postmortem (11, 12), potentially in part due to an increase in available nutrients and energy obtained from fermentation reactions (34). Most *Clostridium* spp. grow strictly in the absence of oxygen and a doubling time of 7.4 minutes (35) which may explain why they so easily colonize the still anaerobic body cavity postmortem. The presence of species of *Lactobacillus*, Enterobacteriaceae, and *Blautia* may be similarly explained. However, the enrichment of *Sediminibacterium* and Rhizobiales in natural deaths and suicides respectively, which are traditionally associated with soil, is harder to understand but may represent colonization by environmental bacteria.

In conclusion, we have identified a number of taxa that may be predictive of manner of death, but this result needs substantial independent validation, and further controlled studies to determine whether the associations are based on biological phenomena.

## Supplemental Material

Figure S1. Linear regression of observed ASV richness by postmortem interval (PMI), faceted by country and organ type.

Figure S2. Linear regression of Shannon Index by postmortem interval (PMI), faceted by country and organ type.

Figure S3. Linear regression of observed ASV richness by body mass index (BMI), faceted by country and organ type.

Figure S4. Linear regression of Shannon Index by postmortem interval (BMI), faceted by country and organ type.

Table S1. Cadaver sampling metadata.

Table S2. Complete ANCOM results for analysis of log2fold change in relative abundance by country.

Table S3. Complete ANCOM results for analysis of log2fold change in relative abundance by organ type, controlling for age, sex, BMI, PMI, ethnicity, and locality.

Table S4. Complete ANCOM results for analysis of log2fold change in relative abundance by manner of death (MOD), controlling for age, sex, BMI, PMI, ethnicity, and locality.

## References

1. Gilbert JA, Quinn RA, Debelius J, Xu ZZ, Morton J, Garg N, Jansson JK, Dorrestein PC, Knight R. 2016. Microbiome-wide association studies link dynamic microbial consortia to disease. Nature 535:94–103.

2. Gilbert JA, Blaser MJ, Caporaso JG, Jansson JK, Lynch SV, Knight R. 2018. Current understanding of the human microbiome. Nat Med 24:392–400.

3. Metcalf CJE, Xu ZZ, Weiss S, Lax S, Van Treuren W, Hyde ER, Song SJ, Amir A, Larsen P, Sangwan N, Haarmann D, Humphrey GC, Ackermann G, Thompson LR, Lauber CL, Bibat A, Nicholas C, Gebert MJ, Petrosino JF, Reed SC, Gilbert JA, Lynne AM, Bucheli SR, Carter DO, Knight R. 2016. Microbial community assembly and metabolic function during mammaliann corpse decomposition. Science 351:158–162.

4. Javan GT FS, Tuomisto S, Hall A, Benbow ME, Mills D.. 2019. An interdisciplinary review of the thanatomicrobiome in human decomposition. Forensic Science, Medicine, and Pathology 15:75–83.

5. Lax S, Hampton-Marcell JT, Gibbons SM, Colares GB, Smith D, Eisen JA, Gilbert JA. 2015. Forensic analysis of the microbiome of phones and shoes. Microbiome 3:21.

6. Fierer N, Lauber CL, Zhou N, McDonald D, Costello EK, Knight R. 2010. Forensic identification using skin bacterial communities. Proc Natl Acad Sci U S A 107:6477–6481.

7. Park J, Kim SJ, Lee J-A, Kim JW, Kim SB. 2017. Microbial forensic analysis of human-associated bacteria inhabiting hand surface. Forensic Science International: Genetics Supplement Series 6:e510–e512.

8. Schmedes SE, Woerner AE, Budowle B. 2017. Forensic human identification using skin microbiomes. Appl Environ Microbiol 83.

9. Kodama WA, Xu Z, Metcalf JL, Song SJ, Harrison N, Knight R, Carter DO, Happy CB. 2018. Trace Evidence Potential in Postmortem Skin Microbiomes: From Death Scene to Morgue. J Forensic Sci doi:10.1111/1556-4029.13949.

10. Burcham ZM HJ, Pechal JL, Krausz KL, Bose JL, Schmidt CJ, Benbow ME, Jordan HR. 2016. Fluorescently labeled bacteria provide insight on postmortem microbial transmigration. Forensic Science International 264:63–69.

11. Javan GT FS, Smith T, Miller J, Wilkinson JE. 2017. Cadaver thanatomicrobiome signatures: the ubiquitous nature of Clostridium species in human decomposition. Front Microbiol 8.

12. Javan GT FS, Can I, Wilkinson JE, Hanson JD, Tarone AM. 2016. Human thanatomicrobiome succession and time since death. Sci Rep 6:195–298.

13. Cobaugh KL SS, DeBruyn JM.. 2015. Functional and structural succession of soil microbial communities below decomposing human cadavers. PLoS ONE 10.

14. Hauther KA CK, Jantz LM, Sparer TE, DeBruyn JM. 2015. Estimating time since death from postmortem human gut microbial communities. J Forensic Sci 60:1234–1240.

15. Pechal JL CT, Benbow ME, Tarone AM, Dowd S, Tomberlin JK.. 2014. The potential use of bacterial community succession in forensics as described by high throughput metagenomic sequencing. Int J Legal Med 128:193–205.

16. Maujean G GT, Fanton L, Malicier D. 2013. The interest of postmortem bacteriology in putrefied bodies. J Forensic Sci 58:1069–1070.

17. Ercolini D, Russo F, Nasi A, Ferranti P, Villani F. 2009. Mesophilic and psychrotrophic bacteria from meat and their spoilage potential in vitro and in beef. Appl Environ Microbiol 75:1990–2001.

18. Deutscher J. 2008. The mechanisms of carbon catabolite repression in bacteria. Curr Opin Microbiol 11:87–93.

19. Rakoff-Nahoum S, Foster KR, Comstock LE. 2016. The evolution of cooperation within the gut microbiota. Nature 533:255–259.

20. Campobasso C, Di Vella G, Introna F. 2001. Factors affecting decomposition and Diptera colonization. Forensic Science International 120:18–27.

21. Amendt J. KR, Zehner R, Bratzke, H.. 2003. Practice of forensic entomology--usability of insect fragments in the estimation of the time of death. Archiv für Kriminologie 214:11–18.

22. Caporaso JG, Lauber, C. L., Walters, W. A., Berg-Lyons, D., Lozupone, C. A., Turnbaugh, P. J., Fierer, N., Knight, R. 2011. Global patterns of 16S rRNA diversity at a depth of millions of sequences per sample. PNAS 108:4516–4522.

23. Caporaso JG, Lauber CL, Walters WA, Berg-Lyons D, Huntley J, Fierer N, Owens SM, Betley J, Fraser L, Bauer M, Gormley N, Gilbert JA, Smith G, Knight R. 2012. Ultra-high-throughput microbial community analysis on the Illumina HiSeq and MiSeq platforms. ISME J 6:1621–1624.

24. Kozich JJ, Westcott, S. L., Baxter, N. T., Highlander, S. K., Schloss, P. D. 2013. Development of a dual-index sequencing strategy and curation pipeline for analyzing amplicon sequence data on teh MiSeq Illumina sequencing platform. Applied and Environmental Microbiology 79:5122–5120.

25. Caporaso JG, Kuczynski, J., Stombaugh, J., Bittinger, K., Bushman, F. D., Costello, E. K., Fierer, N., Gonzalez Peña, A., Goodrich, E. K., Gordon, J. I., Huttley, G. A., Kelley, S. T., Knights, D., Koenig, J. E., Ley, R. E., Lozupone, C. A., McDonald, D., Muegge, B. D., Pirrung, M., Reeder, J., Sevinsky, J. R., Turnbaugh, P. J., Walters, W. A., Widmann, J., Yatsunenko, T., Zaneveld, J., Knight, R. 2010. QIIME allows analysis of high-throughput community sequencing data. Nature Methods 7:335 – 336.

26. Rognes T, Flouri T, Nichols B, Quince C, Mahe F. 2016. VSEARCH: a versatile open source tool for metagenomics. PeerJ 4:e2584.

27. Amir A, McDonald D, Navas-Molina JA, Kopylova E, Morton JT, Zech Xu Z, Kightley EP, Thompson LR, Hyde ER, Gonzalez A, Knight R. 2017. Deblur Rapidly Resolves Single-Nucleotide Community Sequence Patterns. mSystems 2.

28. McMurdie PJ, Holmes S. 2013. phyloseq: an R package for reproducible interactive analysis and graphics of microbiome census data. PLoS One 8:e61217.

29. Oksanen J., Blanchet F. G., Friendly M., Kindt R., Legendre P., McGlinn D., Minchin P. R., O’Hara R. B., Simpson G. L., Solymos P., Henry M., Stevens H., Szoecs E., H. W. 2018. vegan: Community Ecology Package, vR package version 2.5-2. https://CRAN.R-project.org/package=vegan.

30. Wickham H. 2016. ggplot2: Elegant Graphics for Data Analysis. Springer-Verlag, New York, NY.

31. Wickham H, François R, Henry L, Müller K. 2018. dplyr: A Grammar of Data Manipulation, vR package version 0.7.6. https://CRAN.R-project.org/package=dplyr.

32. Hyde ER, Haarmann DP, Lynne AM, Bucheli SR, Petrosino JF. 2013. The living dead: bacterial community structure of a cadaver at the onset and end of the bloat stage of decomposition. PLoS One 8:e77733.

33. Tuomisto S, Karhunen PJ, Vuento R, Aittoniemi J, Pessi T. 2013. Evaluation of postmortem bacterial migration using culturing and real-time quantitative PCR. J Forensic Sci 58:910–916.

34. DeBruyn JM, Hauther KA. 2017. Postmortem succession of gut microbial communities in deceased human subjects. PeerJ 5:e3437.

35. Willardsen RR, Busta—, F. F., and Allen, C. E. 1979. Growth of Clostridium perfringens in three different beef media and fluid thioglycollate medium at static and constantly rising temperatures. J Food Prot 42:144–148.

